# Effect of repeated explicit instructions on visuomotor adaptation and intermanual transfer

**DOI:** 10.1101/2022.03.04.482800

**Authors:** Susen Werner, Heiko K. Strüder

## Abstract

The aim of the present study was to investigate the effect of repeated explicit instructions on visuomotor adaptation, awareness, and intermanual transfer. In a comprehensive study design, 48 participants performed center-out reaching movements before and during exposure to a 60° rotation of visual feedback. Awareness and intermanual transfer were then determined. Twelve participants each were assigned to one of the following adaptation conditions: gradual adaptation, sudden adaptation without instructions, sudden adaptation with a single instruction before adaptation, and sudden adaptation with multiple instructions before and during adaptation. The explicit instructions explained the nature of the visual feedback perturbation and were given using an illustration of a clock face. Analysis of adaptation indices revealed neither increased nor decreased adaptation after repeated instructions compared with a single instruction. Our data also show less adaptation with gradual than with sudden adaptation and less adaptation without than with instruction. These differences were pronounced at the beginning of adaptation; however, by the end of adaptation, all four groups reached similar levels of adaptation. In addition, we found significant group differences for the awareness index, with lower awareness after gradual adaptation than after sudden, instructed adaptation. Intermanual transfer did not differ between groups. However, we found a significant correlation between the awareness and intermanual transfer indices. We conclude that the magnitude of the explicit process cannot be further increased by repeated instruction and that intermanual transfer appears to be largely related to the explicit adaptation process.

## Introduction

Identifying exercise conditions that promote motor learning is critical for the design of optimal movement therapies or sports coaching interventions. One possibility, for example, is to provide learners with information in the form of verbal explanations about the to-be-learned motor task. This form of explicit instruction is discussed in different contexts: in the field of movement therapies (McNevin et al. 2000), injury prevention (Benjaminse and Otten 2010) or sports coaching (Hodges and Franks 2002). And although effective instructions might be crucial for learning a motor skill, there is still disagreement about the role of this form of information (Hodges and Franks 2002). Thus, contradictory effects of explicit instructions on the learning of different motor skills have been described. While some studies show positive effects of instructions on learning (Hardy et al. 1996; Prapavessis and McNair 1999; McNair et al. 2000), other studies reveal a negative effect (Wulf and Weigelt 1997; Gredin and Williams 2016).

In the field of sensorimotor adaptation, on the other hand, the state of research is clearer. In this special form of motor learning, already learned movements are adapted to changed environmental conditions. This plays a role, for instance, in the rehabilitation of stroke patients or in the athletic training of adolescents who have to adapt their motor programs due to length growth. Sensorimotor adaptation is studied, for example, by introducing a perturbation between actual movement and visual feedback during simple target reaching movements. In this case, explicit instructions consist of explanations about the nature of the discrepancy between proprioceptive and visual feedback. They clearly lead to improved adaptation (Benson et al. 2011; Werner et al. 2015), especially early in learning (Taylor et al. 2014; Werner et al. 2015; Neville and Cressman 2018; Wang et al. 2019; Bouchard and Cressman 2021). This improvement in initial adaptation is evident even among older participants, although it is not as pronounced here (Vachon et al. 2020). Explicit instructions also improve visuomotor adaptation in cerebellar patients who normally show significant impairment in motor learning (Taylor et al. 2010) and enable simultaneous adaptation to opposing perturbations (dual adaptation) even under conditions where no learning occurs without instructions (Ayala and Henriques 2021). On the other hand, however, explicit instructions about the nature of the perturbation result in reduced aftereffects (Benson et al. 2011; Werner et al. 2015) and have no influence on hand-localization estimates, thus they do not benefit proprioceptive recalibration (Modchalingam et al. 2019). All these results support the idea that explicit instructions lead to the use of cognitive strategies and thus primarily enhance the explicit process and reduce the implicit process of adaptation (Werner et al. 2015; Neville and Cressman 2018). Accordingly, it is not surprising that instructed participants also show a greater transfer of learning to the untrained hand, as recent research shows that this intermanual transfer can be largely related to the explicit process (Poh et al. 2016; Werner et al. 2019b; Bouchard and Cressman 2021).

In other studies, instead of explicit instructions on the nature of distorted feedback, participants were provided with a specific and effective strategy to cope with the induced visual perturbation. Specifically, they were instructed to counteract the visual distortion by aiming at the adjacent target point. These participants initially show a large reduction in movement errors and good task performance, but their performance deteriorates again over the longer adaptation period (Mazzoni and Krakauer 2006; Taylor et al. 2010; Rand and Rentsch 2015). The cognitive strategy fails because the implicit process of adaptation occurs simultaneously and is added to the unchanging strategic behaviour. This eventually leads to a kind of overlearning. However, this deterioration of performance does not occur if only explicit instructions are given at the beginning of learning instead of a specific strategy. The underlying assumption, which to our knowledge has not been directly investigated, is that here the explicit and implicit processes flexibly come into play. That is, the central nervous system appears to modulate the use of instructions individually and to flexibly adapt the cognitive strategies used to the simultaneous implicit adaptation. However, because instructions have always been given prior to the start of adaptation, it would also be possible for participants to simply “forgot” to use cognitive strategies during the course of adaptation.

In the present study, we therefore investigate the effects of repeated explicit instructions before and during learning on visuomotor adaptation. If we find no overlearning but similar or better adaptation compared to the one-time instruction, the idea of flexible application of explicit and implicit processes is supported. Another goal of our study is to find out whether multiple instructions promote the explicit adaptation process more strongly and thus lead to greater awareness of what is learned and greater intermanual transfer than one-time explicit instructions. In addition, we aim to use a comprehensive study design to confirm previous research findings on the relationship between awareness and intermanual transfer. We will measure adaptation, awareness, intermanual transfer and aftereffects of learning in several conditions: gradual adaptation, sudden adaptation without verbal instructions, sudden adaptation with a one-time instruction before adaptation, and sudden adaptation with several instructions before and during adaptation.

## Method

### Participants

Forty-eight healthy volunteers participated in our study and were randomly divided into four groups of twelve subjects each. All participants were right-handed according to the Edinburgh Handedness Inventory (Oldfield 1971). Participants between the ages of 18 and 30 were recruited and care was taken to ensure that the groups were age-and sex-matched^1^. None of the participants had any prior experience in visuomotor adaptation research. The authors’ local Ethics Committee had approved the procedure of the experiment, all participants gave written informed consent and the experimental protocol was conducted according to the principles expressed in the Declaration of Helsinki.

### Task

The seated participants watched a computer screen through a mirror, such that the virtual image of the screen coincided with the horizontal surface of a digitizing tablet (see Werner, Strüder, & Donchin, 2019). A starting dot appeared for a duration of 3 s plus a random interval of up to 500 ms in the center of the virtual display, and was then replaced by one of eight possible target dots, located 45° apart along an imaginary circle of 5 cm radius about the center. The target dots were displayed for a duration of 1,000 ms. Participants held a digitizing pen in their hand, and reached at each target and back by moving the pen across the digitizing tablet. They were unable to see their arm, due to the mirror and surrounding shrouds; however, pen position was registered and displayed on the screen as a cursor to provide visual feedback about instantaneous hand position. The participants were instructed to reach as quickly and accurately as possible. Reaching movements were performed during episodes of 35 s duration which were interrupted by rest breaks of 5 s. There were about 8 movements per episode.

### Experimental design

All participants were first familiarized with the set-up by performing three episodes under veridical visual feedback, i.e., pen and cursor movements were congruent. Subsequently, participants performed baseline episodes without visual feedback, i.e., no cursor visible, as well as baseline episodes with the left and right hand. In the following adaptation phase of 25 episodes, visuomotor adaptation was induced by rotating the cursor 60 ° CCW around the central point. Group GNI adapted to the gradually introduced perturbation (increase of 3 ° per episode) and groups SNI, SOI, and SSI to the suddenly introduced perturbation. While groups GNI and SNI did not receive instructions on the nature of the perturbation, group SOI was instructed once before the adaptation block and group SSI was instructed several times at the beginning of adaptation and after every 5th episode. Instructions were given with the help of an illustration of a clock face as in Benson et al. (2011). The adaptation phase was followed by two episodes each of inclusion and exclusion in order to test for awareness in a process dissociation procedure as in Jacoby (1991) and as previously applied in visuomotor adaptation research (Werner et al. 2015, 2019b, 2020; Neville and Cressman 2018; Bouchard and Cressman 2021; Maresch et al. 2021; Ayala and Henriques 2021). Before inclusion, participants were instructed to “use what was learned during adaptation”; and before exclusion, participants were asked to “refrain from using what was learned and to perform movements as during baseline”. No visual feedback was given in those episodes and the order of inclusion and exclusion episodes was randomized between participants. During adaptation and awareness test movements were performed with the right hand. In the following test of intermanual transfer, participants used their left hand and there was no visual feedback to avoid confounding transfer with learning benefits to opposite limb learning (Joiner et al. 2013; Poh et al. 2016). Finally, a five-episode washout phase was performed under veridical feedback and with the right hand. Between all individual tests (awareness, intermanual transfer, washout), two refresh episodes were performed under rotated feedback and also using the right hand. Table 1 shows an overview of the experimental protocol.

**Tab. 1:** Experimental protocol.

| **Blockname** | **# of episodes** | **Visual FB** |
| --- | --- | --- |
| Familiarization | 3 | 0 ° |
| Baseline no FB | 2 | - |
| Baseline left hand | 2 | 0 ° |
| Baseline right hand | 2 | 0 ° |
| Adaptation | 25 | G or S<br>60 ° |
| Exclusion/Inclusion | 2 | - |
| Refresh | 2 | 60 ° |
| Inclusion/Exclusion | 2 | - |
| Refresh | 2 | 60 ° |
| Intermanual transfer | 2 | - |
| Refresh | 2 | 60 ° |
| Washout | 5 | 0 ° |

Visual feedback (FB) was either not present (-), veridical (0 °) or rotated (60 °) gradually (G) or suddenly (S). In the gradual condition rotation size was increased in steps of 3 ° per episode. The order of exclusion and inclusion was alternated between participants.

### Data processing

For each movement and each participant, the direction of reaching movement was determined as the angle between the target direction and the line connecting the hand position at motion onset and the position 150 ms later. Motion onset was determined by a velocity threshold of 30 mm/s. From single movement data we calculated mean movement directions for each participant and episode. Data is provided as supporting information S1_Dataset and on OSF (https://osf.io/gau3y/). From this, indices for adaptation, intermanual transfer, and washout were calculated for each subject as

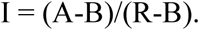

Here, A is the mean value of all movement directions of each block of five episodes for adaptation, both episodes of intermanual transfer or the first two episodes of washout. B is the mean value of all movement directions of both episodes of the baseline condition with the right (adaptation index and washout index) or with the left hand (intermanual transfer) and R is the magnitude of the rotation angle (60°). Furthermore, we determined the exclusion and inclusion indices for each subject as

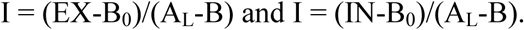

Here, EX and IN are the mean values of all movement directions of both exclusion and inclusion episodes, respectively. B_0_ is the mean value of all movement directions of both baseline episodes without visual feedback and A_L_ is the mean value of all movement directions of the very last adaptation episode. Following the logic of the process dissociation procedure, awareness can be calculated as the difference between exclusion and inclusion performance. Consequently, we determined the awareness index as inclusion index minus exclusion index (Werner et al. 2015, 2019b, 2020). For each index, the value 1 means complete adaptation, awareness, intermanual transfer or washout, while the values −1 to 0 mean no adaptation, awareness, intermanual transfer or washout.

For the statistical analysis baseline and adaptation data were submitted to two analyses of variance (ANOVA) with the between-factor Group (GNI, SNI, SOI, SSI) and the within-factor Block. In addition, we submitted the awareness, intermanual transfer and washout indices to separate one-factor ANOVAs again with the between-factor Group (GNI, SNI, SOI, SSI). Huynh-Feldt adjustments were applied whenever necessary to compensate for heterogeneity of variances. The effect size is reported for significant differences as Eta-squared η2. Significant effects were explored with Fisher LSD post hoc tests. Moreover, partial correlations with the control variable Group (GNI, SNI, SOI, SSI) between awareness index and intermanual transfer or washout index were calculated. The effect size is reported for significant differences as Pearson correlation coefficient R. All these statistical comparisons were performed using SPSS (Version 27.0. Armonk, NY: IBM Corp.).

## Results

### Adaptation

Figure 1 shows the mean movement directions of each episode for all experimental phases and each subject. During the baseline condition, the movement directions of all participants are around 0°, i.e. the movement errors are very small. During the following adaptation phase, group-specific differences become apparent. While in the gradual group the movement direction increases slowly and steadily up to about −50°, the movement directions of the three sudden groups are strongly fluctuating, decrease more suddenly and seem to reach a value of around −50° earlier than in the gradual group. In the following test phases (exclusion and inclusion of the awareness test, intermanual transfer, and washout), the movement directions of the participants in each group are quite similar.

**Fig. 1.**
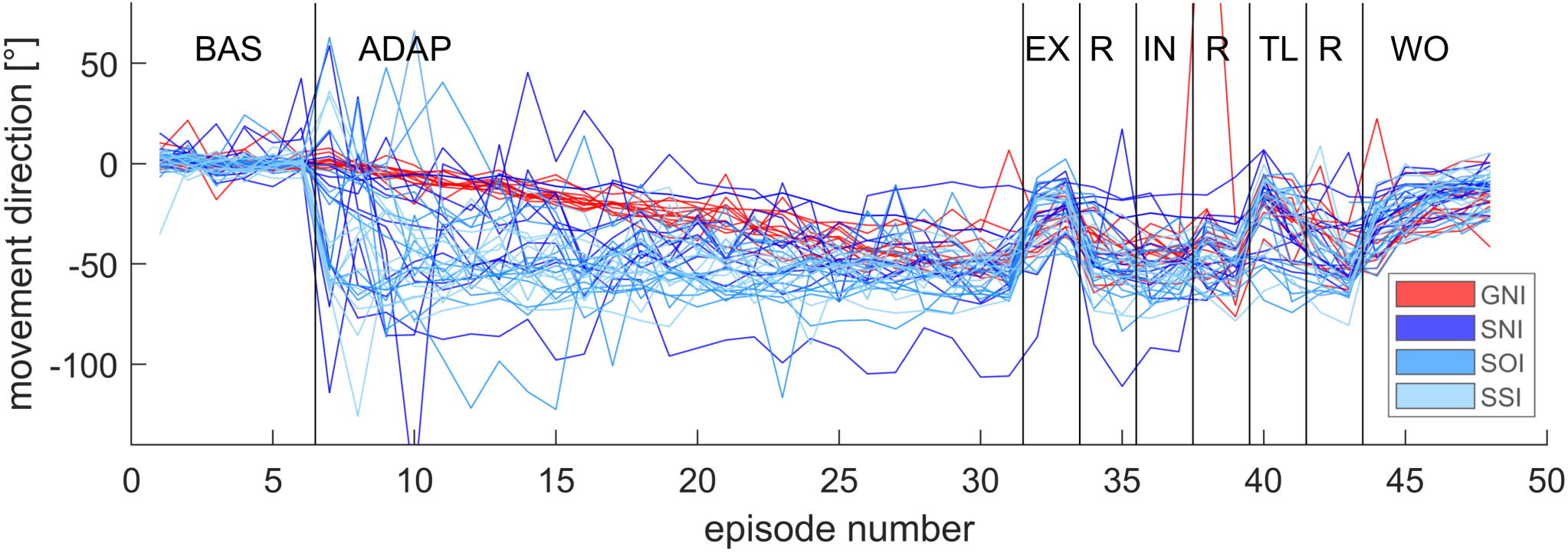
Movement directions of all episodes. Shown are mean movement directions of each episode with respect to target direction of all participants of all groups for baseline (BL), adaptation (ADAP), exclusion (EX), refresh (R), inclusion (IN), transfer to the left hand (TL), and washout (WO) phase. Note that the order of exclusion and inclusion was randomised between participants

Figure 2A depicts the mean group adaptation indices for each block of five episodes. Except for the first block, the course of the two instructed groups SOI and SSI is very similar. The figure also shows that these two groups adapt faster than the sudden group without instructions (SNI). All groups reach an index of about 0.8 at the end of the adaptation phase. It should be noted that the adaptation index measures adaptation as a function of the full rotation magnitude of 60 °. Therefore the adaptation index of GNI increases only slowly over the course of the adaptation phase, although this group shows very small movement errors throughout adaptation. They also reached the full perturbation in the 21st episode, so all groups were exposed to a 60 ° rotation of visual feedback during the 5th block.

**Fig 2.**
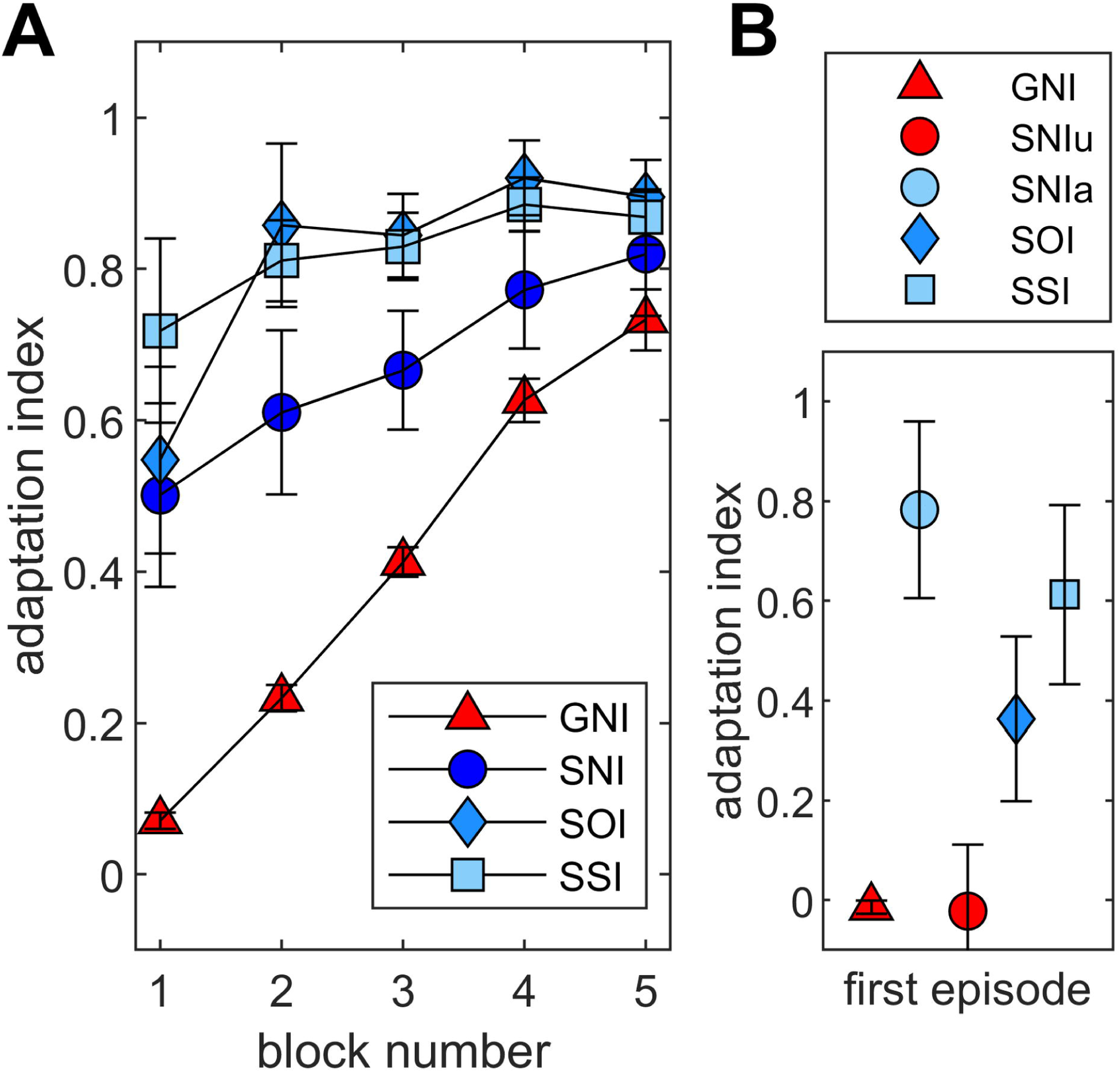
Mean adaptation indices of all blocks and groups. Adaptation index of all blocks for all participants exposed to a gradual rotation (GNI) or a sudden rotation without instructions (SNI), with one-time instruction (SOI) and with several instructions (SSI). Symbols indicate across-subject means and error bars standard deviations **(a)**. In addition, the adaptation indices of these groups for the first episode are depicted. Instead of a mean index for SNI, we show here separate values for unaware (SNIu) and aware (SNIa) participants of this group **(b)**

The above observations of behaviour during baseline episodes and adaptation blocks are confirmed by statistical analysis. ANOVA of the movement directions of baseline phase with the factors Group (GNI, SNI, SOI, SSI) and Block (mean of two episodes) yielded no significant differences between Groups (F(3,44) = 1.404, p = 0.254), Block (F(2,83) = 2.956, p = 0.060), or their interaction (F(6,83) = 0.558, p = 0.753). The ANOVA of the adaptation index, on the other hand, shows significant differences for all comparisons [Group: F(3,44) = 11.644, p < 0.001, *η*^*2*^ = 0.443; Block: F(2,104) = 26.551, p < 0.001, *η*^*2*^ = 0.376 Group × Block: F(7,104) = 2.958, p < 0.01, *η*^*2*^ = 0.168]. The post hoc analysis of the Group effect shows a smaller adaptation index for GNI than for SNI (p < 0.01), SOI (p < 0.001) and SSI (p < 0.001). Mean group values of SNI are smaller than those of SOI and SSI but the analysis does not reach a significant difference [SOI: p = 0.084, SSI: p = 0.066]. Furthermore, the adaptation index does not differ between group SOI and SSI (p = 0.905). The post hoc analysis of the Group × Block interaction shows three things. First, the adaptation index within the groups GNI, SNI, and SOI increases from block 1 to block 5 [GNI: p < 0.001, SNI: p < 0.01, SOI: p < 0.01, SSI: p = 0.175]. This shows that adaptation has occurred. Second, we find no difference between the adaptation indices of all groups for block 5. That is, all groups reach an equal adaptation level at the end of adaptation phase. Third, the individual comparisons show a lower adaptation index for group GNI than all other groups during the first four blocks and a lower index for group SNI than the two instructed groups during blocks 2 and 3 (see Table 2A).

**Tab. 2:** Results of post hoc comparisons for the adaptation index.

| **A** | **Block 1** | **Block 2** | **Block 3** | **Block 4** | **Block 5** | **B** | **Episode 1** |  |
| --- | --- | --- | --- | --- | --- | --- | --- | --- |
|  | GNI | GNI | GNI | GNI | GNI |  | SNIu | SNIa |
| SNI | < **0.01** | < **0.01** | < **0.01** | 0,061 | 0,287 | GNI | 0,977 | < **0.01** |
| SOI | < **0.01** | < **0.001** | < **0.001** | < **0.001** | 0,050 | SOI | 0,154 | 0,121 |
| SSI | < **0.001** | < **0.001** | < **0.001** | < **0.01** | 0,101 | SSI | < **0.05** | 0,524 |
|  | SNI | SNI | SNI | SNI | SNI |  | SNIa |  |
| SOI | 0,768 | < **0.05** | < **0.05** | 0,058 | 0,355 | SNIu | < **0.05** |  |
| SSI | 0,171 | 0,102 | < **0.05** | 0,145 | 0,552 |  |  |  |
|  | SOI | SOI | SOI | SOI | SOI |  |  |  |
| SSI | 0,279 | 0,698 | 0,857 | 0,644 | 0,738 |  |  |  |

Table 2 lists the results of the post hoc comparisons between all groups. Shown are the p values for the significant Group × Block interaction of the repeated measurement ANOVA of the adaptation indices of all blocks (A) and for the significant Group effect of the one-way ANOVA of the adaptation index of the first episode (B).

### Awareness, intermanual transfer and washout

The results of the awareness, intermanual transfer, and washout indices are presented in Figure 3. The figure shows that the mean awareness index of the group GNI is lower than that of SNI, and the latter in turn than SOI and SSI. A similar pattern, but not as pronounced, can be seen for the intermanual transfer index. The washout index data show an opposite pattern with a slightly larger mean washout in GNI than in the other groups. The one-way ANOVA with the factor Group (GNI, SNI, SOI, SSI) yields a significant Group effect for the awareness index (F(3, 47) = 4.076, p < 0.05, *η*^*2*^ = 0.217). Post hoc analysis shows a trend for the difference between GNI and SNI (p = 0.057) and a significantly lower value for GNI than SOI (p < 0.05) and SSI (p < 0.05). Statistical analysis further shows no significant group difference for intermanual transfer (F(3, 47) = 0.899, p = 0.449) and washout indices (F(3, 47) = 1.433, p = 0.246). In addition, partial correlations with the control variable Group yielded a significant correlation between awareness and intermanual transfer index with (R = 0.530, p<0.001) and with washout index (R = −0.445, p = 0.01), respectively.

**Fig 3.**
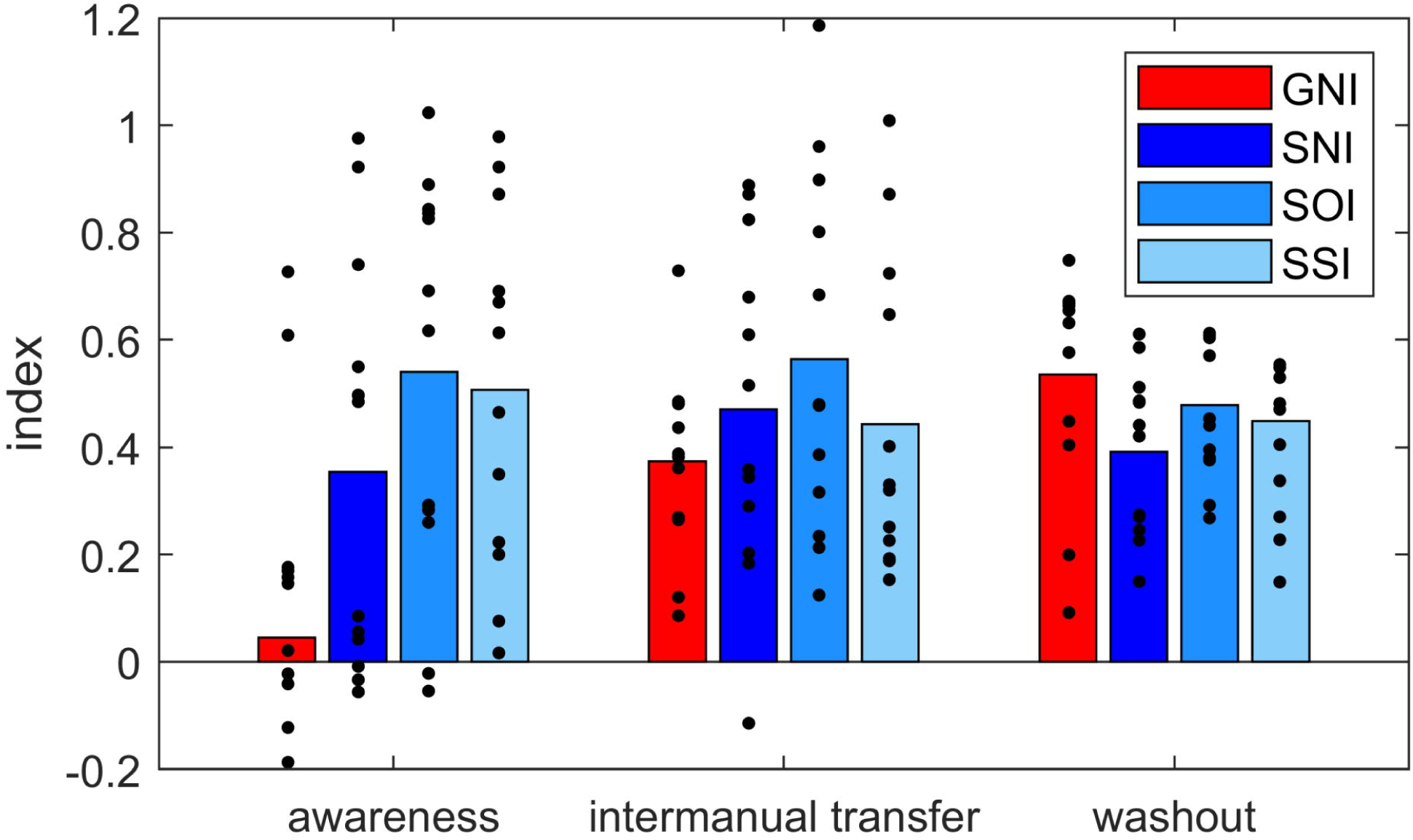
Awareness, intermanual transfer and washout. Shown are group mean values for awareness, intermanual transfer and washout for all participants exposed to a gradual rotation (GNI) or a sudden rotation without instructions (SNI), with one-time instruction (SOI) and with several instructions (SSI). Dots represent individual data

Group SNI adapted to a suddenly introduced perturbation without receiving any instructions. From Figure 3, it can be seen that the mean awareness index of this group is between that of GNI on the one hand and the indices of SOI and SSI on the other. However, it is striking that part of the subjects (n = 6) of SNI show a very low awareness index around the value 0 and another part of the subjects (n = 6) show a high awareness index with values between 0.5 and 1. This indicates that half of the subjects were rather unaware regarding the nature of the perturbation while the other half probably independently inferred the nature of the perturbation due to the large movement error. In addition, previous studies show that instructions lead to reduced movement errors, especially at the beginning of adaptation (Benson et al. 2011; Werner et al. 2015). The comparison of our adaptation data of the groups SNI, SOI and SSI, however, reveals a positive effect of instructions on the adaptation index of block 2 and 3 (see Fig. 2A and Table 2A) but not on the adaptation index of the first block. It is possible that explicit instructions lead to increased early adaptation only if the comparison data come from subjects who are actually unaware concerning the nature of the perturbation. Or, in other words, if uninstructed subjects independently recognize the nature of the perturbation, they might show rapid adaptation just as instructed subjects do.

To find out whether the aware and unaware participants of the SNI group actually differed with respect to their initial adaptation index we divided this group depending on their awareness index and performed an additional one-way ANOVA including the groups GNI, SNI_unaware_, SNI_aware_, SOI and SSI on the adaptation index of the first episode. Indeed, this analysis yields a significant group difference (F(4, 47) = 3.879, p < 0.01, *η*^*2*^ = 0.265). As can be seen in Figure 2B and from Table 2B, the adaptation index of SNI_unaware_ is lower than that of SNI_aware_. Moreover, we find significant differences between SNI_unaware_ and SSI and between SNI_aware_ and GNI, respectively. This implies that those participants who independently gained awareness regarding the nature of the perturbation have a similar advantage at the beginning of adaptation as instructed participants. We do not find this effect on the intermanual transfer and washout indices. Here, the statistical analyses do not reveal any significant group differences [intermanual transfer: F(4, 47) = 1.043, p = 0.396; washout: F(4, 47) = 1.565, p = 0.201].

## Discussion

The aim of the present study was to determine the effect of multiple explicit instructions on visuomotor adaptation, awareness, and intermanual transfer of learning. In a broad study design, 48 participants adapted to a 60° rotation of visual feedback. Twelve participants were each assigned to one of the following conditions: gradual adaptation (GNI), sudden adaptation without instructions (SNI), sudden adaptation with a one-time instruction (SOI) before adaptation, and sudden adaptation with several instructions (SSI) before and during adaptation. The explicit instructions explained the nature of the perturbation of visual feedback and were given with the help of an illustration of a clock face (Benson et al. 2011).

### Multiple versus one-time instructions

Our results show no improvement in adaptation with multiple explicit instructions compared to a one-time instruction prior to the onset of learning. Thus, the effect of explicit instructions is not enhanced by repetition. Accordingly, while it is common in sports training to repeat instructions, this does not seem to provide a learning advantage in sensorimotor adaptation. On the other hand, we also find no deterioration in adaptation from multiple instructions, as is the case when using a specific cognitive strategy (Mazzoni and Krakauer 2006; Taylor et al. 2010; Rand and Rentsch 2015). Moreover, we find no difference in awareness or intermanual transfer between the two groups SOI and SSI. In summary, these results can be interpreted as follows: First, multiple instructions do not seem to further increase the size of the explicit process. We would otherwise need to find greater awareness in SSI than SOI (Bouchard and Cressman 2021). Second, we find no evidence that the one-time instruction was “forgotten”. We would otherwise have to find lower awareness in SOI than SSI. This result is also consistent with a previous finding that the size of the explicit process after a single explicit instruction remains the same throughout the adaptation phase (Neville and Cressman 2018). Third, the data suggest that the central nervous system applies cognitive strategies flexibly and accounts for concurrent implicit adaptation. We would otherwise have to find a deterioration of adaptation with multiple instructions (Mazzoni and Krakauer 2006; Taylor et al. 2010; Rand and Rentsch 2015).

### Instructions versus no instructions

Our data reveal a trend for greater general adaptation among instructed participants (SOI and SSI) than among uninstructed participants (SNI). Examination of the individual adaptation blocks shows that this difference is more pronounced at the beginning of the adaptation phase. This is consistent with the results of previous studies (Benson et al. 2011; Taylor et al. 2014; Werner et al. 2015; Neville and Cressman 2018; Wang et al. 2019; Bouchard and Cressman 2021). In the present experiment, we provided explicit explanations about the nature of the visual perturbation without examining the resulting consequences for the participants in more detail. However, previous studies show enlarged reaction times (Benson et al. 2011) and prefrontal cortex involvement (Anguera et al. 2010) in instructed subjects. Taken together, these results suggest that the instructed participants used more cognitive strategies. This may also be true for some participants in the non-instructed (SNI) group. This is because our data reveal, for the first time, that those participants who independently develop an awareness of the nature of the perturbation show exactly the same improvements in adaptation as the instructed participants. Similar results were also found in an experiment on visuomotor sequence learning: Participants who discovered the rules spontaneously showed similar behaviour to participants who were instructed. Both showed fewer errors in a transfer session than unaware participants (Tanaka and Watanabe 2017). In previous studies, explicit instruction about the nature of the perturbation did lead to improved adaptation, but this was at the expense of reduced aftereffects of visuomotor (Benson et al., 2011; Werner et al., 2015) or locomotor learning (French, Morton, Charalambous, & Reisman, 2018). Accordingly, while we find no group differences for the washout index between instructed and non-instructed participants, we do find a negative correlation between the amount of awareness gained during adaptation and aftereffect size.

### Intermanual transfer

Another aim of the present study was to examine the relationship between awareness and intermanual transfer in a comprehensive study design. Consistent with previous research we found greater intermanual transfer in more aware participants across all groups. Our data thus confirm the idea that transfer of learning to the other hand is largely related to the explicit process of adaptation (Poh et al. 2016; Werner et al. 2019b; Bouchard and Cressman 2021). This is also consistent with recent research showing that the fast adaptation process (Smith et al. 2006) previously associated with the explicit process (McDougle et al. 2015) is responsible for the generalization of learning (Xing and Saunders 2021). In the more detailed analysis of the individual groups, we find a difference in awareness between the gradual and the sudden (especially the instructed) groups, but we do not find a group effect of intermanual transfer. A possible explanation arises from the results of our previous study, in which we showed that a different intermanual transfer after gradual and sudden adaptation occurs only when adapting to a larger rotation (e.g., 75°, Werner et al. 2019a). This is because during adaptation to smaller rotation angles (e.g., 30°), participants in the sudden group are just as unaware as those in the gradual group. This could also be the case for a 60° rotation angle. However, the awareness of group SNI in the present study is clearly pronounced with a mean awareness index of 0.36 and is even greater than after sudden adaptation to a 75° rotation with an index of about 0.3 (Werner et al. 2019b). An alternative explanation is based on the fact that the amount of transfer cannot be fully explained by awareness of what is learned. There seems to be both an implicit and an explicit component of intermanual transfer (Poh et al. 2016; Werner et al. 2021). In our data, we cannot distinguish these two components. It is possible that in the GNI group the proportion of implicit intermanual transfer is particularly pronounced, for example, due to martial arts training, greater general athletic activity (Werner et al. 2021) or stronger right-handedness (Lefumat et al. 2015; Werner et al. 2021).

## Conclusion

Our results show neither an improvement nor a deterioration of adaptation when multiple explicit instructions are given compared to a single instruction before learning begins. We conclude that the size of the explicit process may not further be increased by instructions and that the central nervous system can flexibly apply cognitive strategies and account for simultaneous implicit adaptation. In addition, we find a positive relationship between awareness of what has been learned and intermanual transfer. The transfer of learning to the other hand thus appears to be largely related to the explicit adaptation process.

## Acknowledgments

This work was supported by a scholarship of the German Sport University for female young scientists awarded to SW. Thanks are due to Christian Rickers for data collection.

## Data accessibility statement

All data are available in the online repository (https://osf.io/gau3y/)

## Supporting information caption

**S1_Dataset:** Mean movement directions of each episode (baseline, adaptation, awareness test, refresh, intermanual transfer, and washout) of each participant and all groups (GNI, SNI, SOI, SSI).

## Footnotes

1 The handedness questionnaires as well as the forms with the personal data of the participants were unfortunately destroyed in the course of a move to a new institute building. Therefore, it is not possible for us to give the exact number of men and women as well as the exact age of the participants.

